# Forelimb movements evoked by optogenetic stimulation of the macaque motor cortex

**DOI:** 10.1101/2019.12.13.876219

**Authors:** Hidenori Watanabe, Hiromi Sano, Satomi Chiken, Kenta Kobayashi, Yuko Fukata, Masaki Fukata, Hajime Mushiake, Atsushi Nambu

## Abstract

Optogenetics has become an indispensable tool for investigating brain functions. Although non-human primates are particularly useful models for understanding the functions and dysfunctions of the human brain, application of optogenetics to non-human primates is still limited. In the present study, we generated an effective adeno-associated viral vector serotype DJ to express channelrhodopsin-2 (ChR2) under the control of a strong ubiquitous CAG promoter and injected into the somatotopically identified forelimb region of the primary motor cortex in macaque monkeys. ChR2 was strongly expressed around the injection sites, and optogenetic intracortical microstimulation (oICMS) through a homemade optrode induced prominent cortical activity: Even single-pulse, short duration oICMS evoked long-lasting repetitive firings of cortical neurons. In addition, oICMS elicited distinct forelimb movements and muscle activity, which were comparable to those elicited by conventional electrical ICMS. The present study removed obstacles to optogenetic manipulation of neuronal activity and behaviors in non-human primates.

---

Optogenetics is a technique to manipulate neuronal excitability by using genetically coded, light-gated ion channels or pumps (opsins) and light. Optogenetics is widely used and has become an indispensable tool for investigation of the functions of the nervous system^1, 2, 34^. This method has several advantages over classical tools, such as electrical stimulation and pharmacological blockade, because it can excite or inhibit a specific population of neurons with a high time resolution. Optogenetics has been successfully used to modulate behaviors in rodents^5, 6, 7^. However, its application in non-human primates is still rather limited. Optogenetic stimulation has been attempted to induce and/or modulate body and eye movements, which can be easily elicited by electrical intracortical microstimulation (eICMS) with weak currents. Eye movements can be successfully modulated by optogenetic activation or inactivation of the cerebral cortex ^8, 9, 10, 11, 12^. On the other hand, optogenetic stimulation of the motor cortices in monkeys failed to induce apparent body movements, although it activated or modulated cortical activity^13, 14, 15^. This may be because opsins were not sufficiently expressed in neurons, and/or lights did not effectively penetrate the monkey brain tissue^13, 16^. Inducing movements by optogenetics is a very important next step in non-human primate research. In the present study, we have overcome these drawbacks by injecting effective viral vectors into the identified primary motor cortex (M1) and by using effective delivery of stronger light through optrodes that combine an optical fiber with recording and electrical stimulating electrodes. After these modifications, optogenetic intracortical microstimulation (oICMS) of the M1 successfully induced forelimb movements and muscle activity that were comparable to those induced by eICMS. It also allowed us to record neuronal activity evoked by oICMS and compare the effects of oICMS and eICMS.

## Results

### Gene transfer mediated by adeno-associated virus (AAV) vectors

To achieve the effective AAV vector-mediated gene transfer in the macaque brain, we compared different serotypes of AAV vectors, i.e., 2, 5, and DJ. First, electrophysiological mapping was carried out in the M1 of a macaque to determine the sites for viral vector injections (Supplementary Fig. 1a). Then, we injected each serotype of AAV vectors containing the ubiquitous CAG promoter and the enhanced green fluorescent protein (EGFP) transgene (1.5×10^9^ viral genome (vg)/μl, 1 µl/site, 2-5 sites) into the M1 of each monkey (Supplementary Table 1). Two to four weeks after AAV injection, when the transgene was expected to be expressed, we processed the brain sections and observed EGFP under a fluorescent microscope with the same exposure for the three different serotypes. The fluorescence signals were most conspicuous around the injection sites of the AAV-DJ vector (Supplementary Fig. 1b). The areas and number of cells transduced by the AAV-DJ vector were larger than those transduced by AAV2 and AAV5 (Supplementary Fig. 1c). The intensity of fluorescence signals at the single-neuron level by the AAV-DJ vector was higher than that by AAV2 and AAV5 (Supplementary Fig. 2a). Observing expression of NeuN, parvalbumin (PV), and glial fibrillary acidic protein (GFAP) among EGFP positive cells (Supplementary Fig. 2b, c) showed that the major cell type expressing EGFP was NeuN-positive neurons, and that glial cells expressing EGFP were a small population (Supplementary Fig. 2b). The AAV-DJ vector induced EGFP expression at terminals in the putamen and at axons presumably in the lateral corticospinal tract of the spinal cord, but no retrogradely labeled cells were found in the thalamus (Supplementary Fig. 2d). In addition, the EGFP protein level induced by the AAV-DJ vector in the mouse brain was higher than that induced by AAV2 and AAV5 (Supplementary Fig. 3).

Therefore, we chose the AAV-DJ vector and generated a vector containing channelrhodopsin-2 (ChR2) with a point mutation [hChR2(H134R)] and a strong ubiquitous promoter CAG, i.e., AAV-DJ-CAG-hChR2(H134R)/EYFP or AAV-DJ-CAG-hChR2(H134R)/tdTomato. We then injected the either one of them into 8-15 sites (1 µl/site) in layer 5 of the forelimb region of the M1 (Fig. 1a, b; Supplementary Table 1) based on electrophysiological mapping. Three weeks after AAV injection when ChR2 was expected to be expressed, recording and stimulating experiments were started. Postmortem histological examinations showed distinct fluorescent signals of hChR2(H134R)/EYFP or hChR2(H134R)/tdTomato around the injection sites (Fig. 1c): The area covered by hChR2(H134R)/EYFP signals (72.0 mm^2^, Monkey NR, Left M1) was larger than that by the tdTomato-containing vector (16.7 mm^2^, Monkey NR, Right M1).

**Fig. 1.**
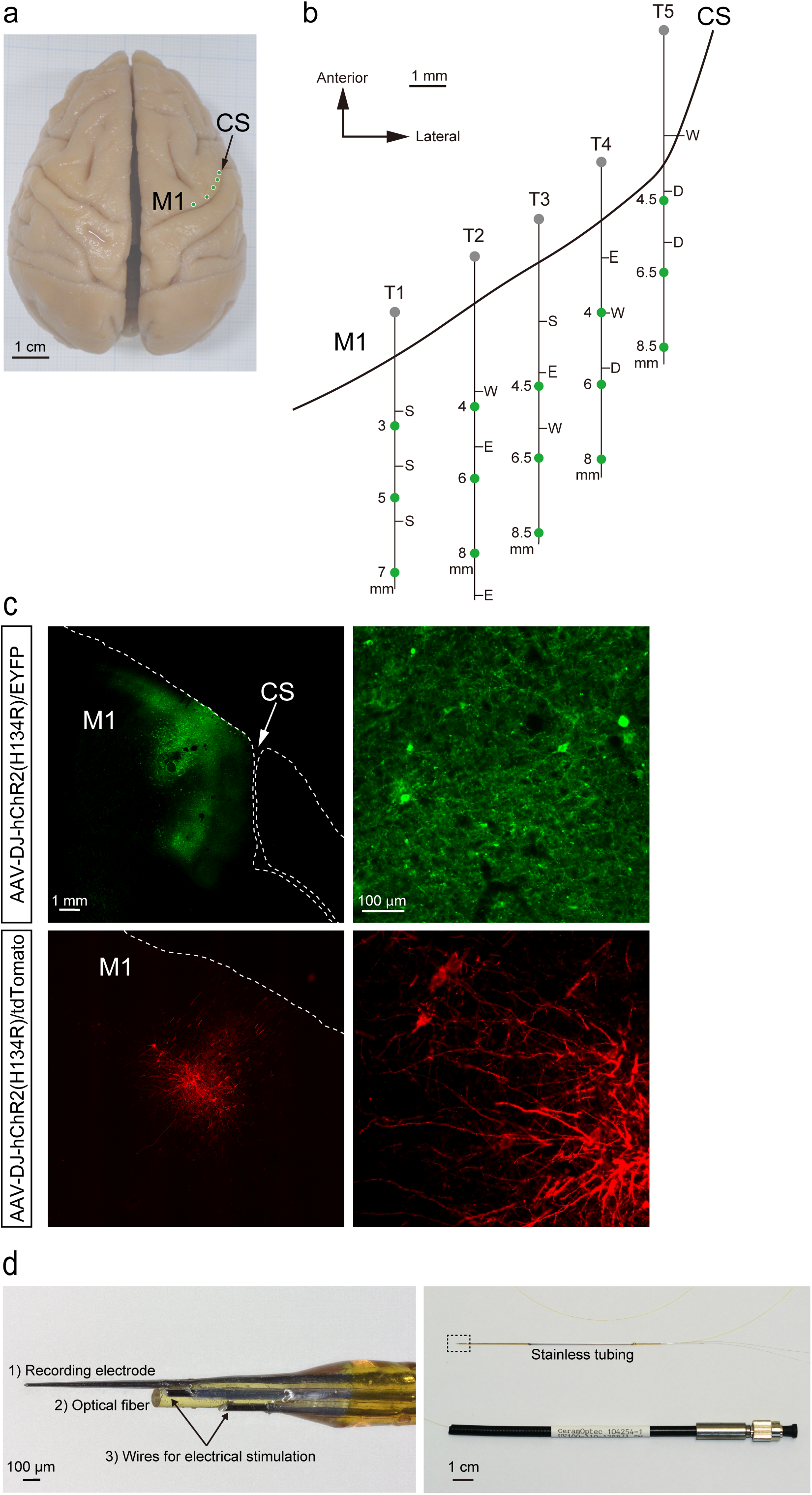
Mapping of the motor cortex and injection of viral vectors. **a** Top view of the whole brain of a macaque (Monkey HJ). Green circles indicate penetration sites for AAV injections. CS, central sulcus; M1, primary motor cortex. **b** Electrophysiological mapping of the right M1. Somatotopic arrangements in the anterior bank of the CS are shown along with depths from the cortical surface. Each letter indicates a somatotopic body part: D, digit; E, elbow; S, shoulder; W, wrist. The AAV vector was injected in the sites indicated by the green circles. **c** Expression of hChR2(H134R)/EYFP and of hChR2(H134R)/tdTomato mediated by AAV-DJ vectors in the M1 (Monkey NR). **d** Photograph of the tip (*left*) and whole view (*right*) of the optrode. 1) A glass coated tungsten electrode for recording, 2) an optical fiber for optical stimulation, and 3) a pair of tungsten wires for bipolar electrical stimulation were bundled and inserted into polyimide tubing (*left*). The base of the optorode was covered with stainless tubing (*right)*. The other end of the optical fiber was jacketed with a protective tubing with a length of around 15 cm and had a fiber-optic connector. The rectangular area in the right photograph is enlarged in the left.

### Neuronal activity in the M1 evoked by optogenetic stimulation

We made an optrode by bundling 1) a glass coated tungsten electrode for recording, 2) an optical fiber for optical stimulation, and 3) a pair of tungsten wires for bipolar electrical stimulation (Fig.1d, see Methods for details). This optrode enabled us to record neuronal activity induced by optical and electrical stimulation. To examine responses of cortical neurons to optical stimulation, we inserted the optrode into the forelimb regions of the M1, where the AAV-hChR2 vector was injected (Supplementary Table 1). Then, oICMS was delivered through the optical fiber, which was connected to a solid-state laser source (473 nm blue laser), and neuronal activity was recorded through the recording electrode. While eICMS usually produced large stimulus artifacts and hindered recording of neuronal responses just after stimulation, oICMS induced negligible stimulus artifacts and therefore enabled us to analyze evoked cortical activity precisely.

Both single-pulse and repetitive oICMS within 1 mm from the injection sites of the viral vector successfully induced neuronal activity in all five monkeys examined (Fig. 2, Supplementary Table 2). Figure 2a shows a typical example of recordings from a single neuron. Low-intensity oICMS less than 1 mW (corresponding to 127 mW/mm^2^) was sufficient to evoke spikes in cortical neurons (Fig. 2a, 0.4mW corresponding to 51 mW/mm^2^), suggesting a high level of ChR2 expression. The minimum intensity and duration required to evoke cortical activity were 0.4 mW and 0.1 ms, respectively. Such low-intensity stimulation just above the threshold evoked an action potential at a latency of 2-4 ms (Fig. 2a, 0.4 mW) with time jitter. A slightly higher-intensity oICMS evoked an action potential with a short and constant latency (less than 1.5 ms) (* in Fig. 2a, 0.9 mW corresponding to 115 mW/mm^2^, 1 ms) followed by a spike with time jitter. Cortical stimulation usually induces both directly evoked excitation and synaptically induced excitation via axon collaterals of cortical neurons^17, 18, 19^. Action potentials at a short and constant latency and the following spikes with time jitter may correspond to directly evoked and synaptically induced excitation, respectively. Stronger oICMS induced repetitive directly evoked spikes (* in the bottom trace of Fig. 2a, 0.9 mW, 2 ms). Even higher-intensity oICMS often induced a large deflection from the baseline (arrowheads in Fig. 2a, 3.6 mW, 15 mW, corresponding to 458 and 1910 mW/mm^2^). The deflection is unlikely an artifact, because (1) it was not induced by a 589 nm yellow laser light, and (2) it was not observed at non-injection sites (Supplementary Fig. 4). Instead, the deflection seems to be composed of population spikes of multiple neurons and/or local field potentials that occurred at the same time. The deflection started just after the onset of oICMS, corresponding to an extremely fast rising time of ChR2-induced photocurrents with no visible delay^20^. The latency of deflection peaks was typically <1.0 ms, and more intense stimulation shortened the latency (Fig. 2a, 3.6 mW, 15 mW). The deflection was immediately followed by repetitive action potentials with a small amplitude that gradually recovered (arrows in Fig. 2a, 3.6 mW, 15 mW). This phenomenon is interpreted that high-intensity oICMS induces large depolarization in cortical neurons, which lasts long after the cession of laser illumination (up to 10 ms), and truncates the shapes of action potentials because of depolarization block^21^. These complex responses evoked by oICMS were also observed in eICMS (Supplementary Fig. 5a). Both oICMS and eICMS induced excitation at a distance of 2.2 mm from the stimulation site (Supplementary Fig. 5b), suggesting that they both transsynaptically activated neurons in certain areas.

**Fig. 2.**
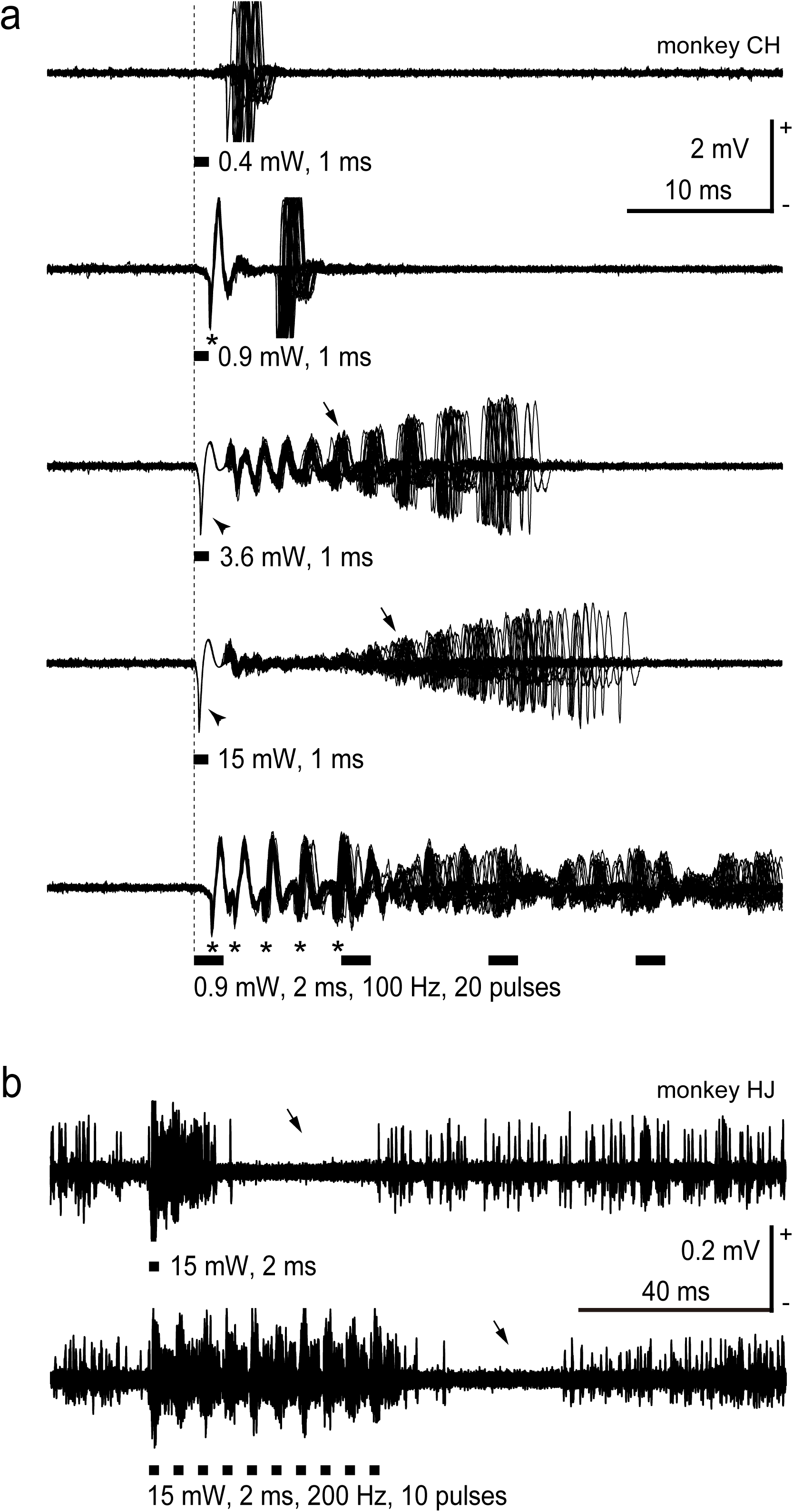
Raw traces of M1 neuronal activity evoked by optogenetic intracortical microstimulation (oICMS). **a** An example of M1 neuronal activity induced by various intensities of single-pulse (0.4, 0.9, 3.6, 15 mW strength corresponding to 51, 115, 458, and 1910 mW/mm^2^, respectively, 1 ms pulse duration) and repetitive (0.9 mW strength corresponding to 115 mW/mm^2^, 2 ms pulse duration, 100 Hz, 20 pulses) oICMS. The neuronal activity from one and the same neuron probably located in layer 5 was recorded, and thirty traces are overlaid. A vertical dashed line and thick black horizontal lines indicate the beginning and duration of laser illumination, respectively. *, action potentials at a short and constant latency that were most likely evoked directly; arrowheads, deflections from the baseline that were composed of population spikes of multiple neurons and/or local field potentials occurring at the same time; arrows, deteriorated action potentials that were truncated due to long-lasting depolarization. Note that no spikes were observed before stimulation, suggesting that the recording neuron was not injured by the optrode. **b** Another example of M1 neuronal activity induced by single-pulse (15 mW strength corresponding to 1910 mW/mm^2^, 2 ms pulse duration) and repetitive (15 mW strength, 2 ms pulse duration, 200 Hz, 10 pulses) oICMS shown on a long time scale. Multi-unit activity was recorded probably in layer 5. Excitation induced by oICMS was followed by inhibition (arrows).

Figure 2b shows a typical example of multiunit activity. The excitation evoked by single-pulse and repetitive oICMS was commonly followed by inhibition of firings (arrows in Fig. 2b; 198/359 recording neurons, 55%). The inhibition may be caused by inhibitory cortical interneurons expressing ChR2 under the control of a ubiquitous CAG promoter and/or medium to slow afterhyperpolarization following high-frequency repetitive firings^22^. Figure 2b also shows that action potentials followed each stimulus pulse at 200 Hz (Fig. 2b, 15 mW corresponding to 1910 mW/mm^2^, 2 ms, 200 Hz, 10 pulses).

### Forelimb movements evoked by optogenetic stimulation

Next, we examined whether the oICMS evoked M1 activity observed above could induce body movements. The tip of the optrode was placed in the forelimb region of the M1 where large neuronal responses were evoked by oICMS. Repetitive oICMS (1.8-15 mW, corresponding to 229-1910 mW/mm^2^, 1-2 ms pulse duration, 100-333 Hz, 10-20 pulses) clearly induced short-duration phasic movements in the contralateral forelimb of all five monkeys examined (Supplementary Table 2). Supplementary movie 1 shows such examples: finger extension, muscle twitching in the forearm, and shoulder elevation. These movements were similar to those evoked by conventional eICMS. oICMS and eICMS shared the following features: (1) They elicited movements in the same parts of the forelimb in concert with the somatotopic map obtained in M1 mapping for vector injection, (2) Single-pulse oICMS/eICMS induced movements, and repetitive oICMS/eICMS was more effective, (3) The amplitude of movements evoked by oICMS was comparable to that evoked by eICMS, and (4) Low-intensity oICMS/eICMS just above the motor threshold typically evoked confined movements, such as single-joint movements or muscle twitches, whereas higher-intensity oICMS/eICMS induced large movements of multiple joints and body parts.

### Muscle activity evoked by optogenetic stimulation

To examine the relationship between M1 activity and body movements, we simultaneously recorded M1 activity by the optrode and electromyogram (EMG) of the forelimb with surface dish electrodes in four monkeys (Supplementary Table 2). Both single-pulse and repetitive oICMS induced M1 activity (Fig. 3a, b, top traces), and shapes of spikes during oICMS were truncated by strong depolarization as seen in Fig. 2a. They also induced muscle activity of extensor and flexor muscles (Fig. 3a, b, bottom two traces). The mean threshold to elicit muscle activity by single-pulse oICMS (10 ms duration) was 10.5±6.3 mW (corresponding to 1337±802 mW/mm^2^) (range, 0.4-15 mW corresponding to 51-1910 mW/mm^2^). Higher intensity of oICMS was more effective than lower intensity as in the case of eICMS. Repetitive oICMS evoked stronger M1 and muscle activity than single-pulse oICMS (Fig. 3; Fig. 4a, *left*) as in the case of repetitive eICMS (Fig. 4a, *right*). The latency of muscle activity evoked by oICMS was 14.0±3.9 ms (n=52 sites; 15 mW corresponding to 1910 mW/mm^2^, 2ms, 200 Hz, 5 pulses), which was comparable to that induced by eICMS (12.8±3.2 ms, n=39 sites; 40-65 µA, 0.2ms, 333 Hz, 5 pulses).

**Fig. 3.**
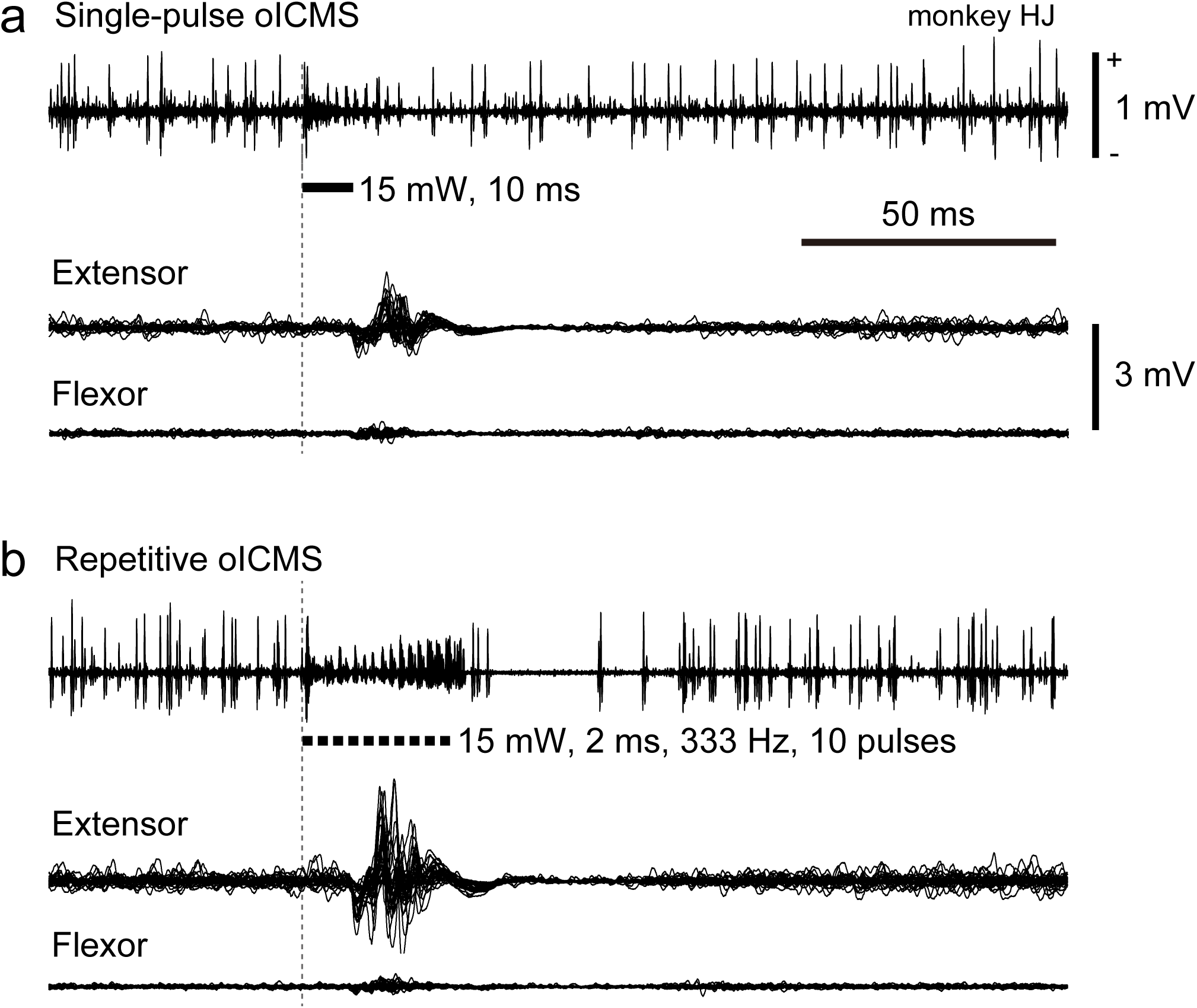
Simultaneously recorded M1 and muscle activities evoked by oICMS. **a, b** M1 activity (top trace) and electromyograms (EMG; bottom two traces) induced by single-pulse (**a**, 15 mW strength corresponding to 1910 mW/mm^2^, 10 ms pulse duration) and repetitive (**b**, 15 mW strength, 2 ms pulse duration, 333 Hz, 10 pulses) oICMS. M1 neuronal activity was recorded probably from neurons in layer 5. EMG was recorded from the extensor carpi radialis longus (Extensor) and flexor carpi radialis (Flexor) muscles. Vertical dashed lines and thick black horizontal lines indicate the beginning and duration of laser illumination, respectively. Calibration in **a** is applicable to **b**.

**Fig. 4.**
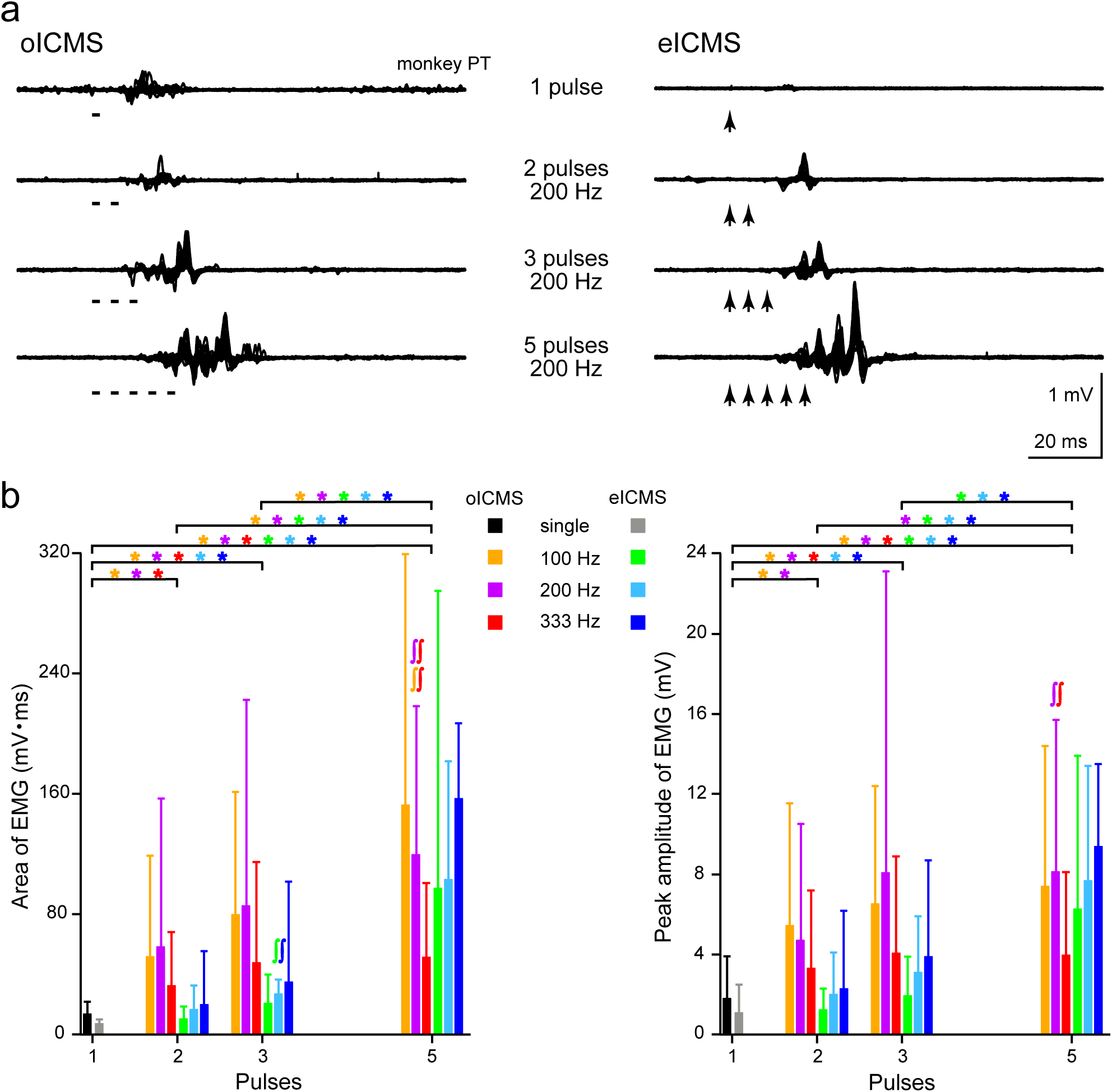
Muscle activity evoked by oICMS and electrical ICMS (eICMS). **a**, Raw EMG recorded from the flexor carpi radialis muscle. One, two, three, and five stimulus pulses of oICMS (*left*, 15 mW strength corresponding to 1910 mW/mm^2^, 2 ms pulse duration, 200 Hz) and eICMS (*right*, 40-65 µA strength, 0.2 ms pulse duration, 200 Hz) were applied to the same site. **b** The area (*left*) and peak amplitudes (*right*) of EMG evoked by oICMS or eICMS with one, two, three, and five pulses at different frequencies. Both oICMS and eICMS were applied, and the EMG data were averaged across stimulating sites (33 and 19 sites for oICMS and 19 and 20 sites for eICMS in Monkeys HJ and PT, respectively). Bar graphs represent means and SDs. Different colors indicate different frequencies (single pulse, 100, 200, and 333 Hz) of oICMS (warm colors) and of eICMS (cold colors). * *p*<0.05, significantly different between stimulus pulses in the frequency indicated by the color; ∫∫ *p*<0.05, significantly different from each other between stimulus frequency indicated by the color (Friedman test with Dunn’s post hoc test).

To examine the effects of stimulus pulse numbers and frequencies on muscle activity, we compared the area and peak amplitude of EMG evoked by oICMS and eICMS with different numbers of pulses (one, two, three, and five pulses) and frequencies (100, 200, and 333 Hz) (52 sites for oICMS, 39 sites for eICMS; Fig. 4b). Five pulses of oICMS and eICMS showed comparable areas and peak amplitudes of EMG (Fig. 4b). When the number of stimulus pulses was decreased, the area and peak amplitude by oICMS were decreased more slowly than those by eICMS. Regarding the area of EMG (Fig. 4b, *left*), two and three pulses of oICMS at 100 and 200 Hz induced significantly smaller EMG than five pulses of oICMS (Friedman test with Dunn’s post hoc test) (for two pulses, *p*<0.0001 at 100 and 200 Hz; for three pulses, *p*=0.0002 at 100 Hz, *p*=0.013 at 200 Hz), but no significant differences at 333 Hz. However, the peak amplitude of EMG (Fig. 4b, *right*) by two, three, and five pulses of oICMS was similar to each other (*p*>0.05), except that the peak amplitude by two pulses of oICMS was significantly smaller than that by five pulses at 200 Hz (*p*<0.0001). One pulse of oICMS induced significantly smaller EMG than two, three, or five pulses of oICMS in both the area and peak amplitude (area; for two pulses, *p*<0.0001 at 100 and 200 Hz, *p*=0.009 at 333 Hz; for three pulses, *p*<0.0001 at 100 and 200 Hz, *p*=0.0002 at 333 Hz; for five pulses, *p*<0.0001 at 100, 200, and 333 Hz) (peak amplitude; for two pulses, *p*=0.0003 at 100 Hz, *p*=0.005 at 200 Hz; for three pulses, *p*<0.0001 at 100 and 200 Hz, *p*=0.005 at 333 Hz; for five pulses, *p*<0.0001 at 100 and 200 Hz, *p*=0.002 at 333 Hz), except the peak amplitude by one and two pulses at 333 Hz was similar to each other (*p*>0.05). On the other hand, both the area (Fig. 4b, *left*) and peak amplitude (Fig. 4b, *right*) by one, two, and three pulses of eICMS were all significantly smaller than those by five pulses of eICMS at 100, 200, or 333 Hz (area; for one and two pulses, *p*<0.0001 at 100, 200, and 333 Hz; for three pulses, *p*<0.0001 at 100 and 200 Hz, *p*=0.025 at 333 Hz) (peak amplitude; for one and two pulses, *p*<0.0001 at 100, 200, and 333 Hz; for three pulses, *p*<0.0001 at 100 Hz, *p*=0.0007 at 200 Hz, *p*=0.004 at 333 Hz). The area and peak amplitude by one and two pulses of eICMS were similar to each other (*p*>0.05), and those by two and three pulses of eICMS were also similar to each other (*p*>0.05). These results suggest that oICMS induces muscle activity more effectively than eICMS at fewer pulses. Concerning the frequency of repetitive stimulation, 333 Hz-stimulation was less effective in oICMS: The area of EMG by 333 Hz-oICMS was significantly smaller than those by 100 and 200 Hz-oICMS (*p<*0.0001 for 100 Hz and 200 Hz) (Fig. 4b, *left*), and also the peak amplitude by 333 Hz-oICMS was significantly smaller than that by 200 Hz-oICMS (*p<*0.0001) (Fig. 4b, *right*), suggesting that the effects of oICMS were saturated around 300 Hz. On the other hand, the effects of different frequencies were smaller in eICMS: 100, 200, and 333 Hz-eICMS evoked similar areas and peak amplitudes of EMG (Fig. 4b), except that three pulses of eICMS at 100 Hz induced smaller area than that at 333 Hz (*p*=0.018).

## Discussion

Introduction of optogenetics has revolutionized stimulation methods^1, 2, 3, 4^ and provided several advantages over electrical stimulation. Optogenetics can selectively stimulate a specific population of neurons or specific pathways by labeling them with light-sensitive opsins using transgenic techniques and/or viral vectors. So far, most optogenetic studies have been performed in rodents and reported photo-induced neuronal and behavioral changes. In the early optogenetic studies, optical stimulation of the motor cortex in rodents expressing ChR2 induced apparent whisker deflection or limb movements^6, 7, 23, 24^. Despite these remarkable results in rodents, optogenetic stimulation of the sensorimotor cortex in macaque monkeys using ChR2 failed to induce arm and hand movements^13, 14^. In another study, although gamma oscillations were induced in the macaque motor cortex by optogenetic stimulation using a ChR2 variant, no clear behavioral modulation was observed^15^. Transduction of enhanced NpHR (eNpHR) into the macaque M1 hand-arm region caused suppression of M1 neuronal activity, but no obvious modulation of reaching and grasping behaviors was found^13, 25^. On the other hand, optogenetic stimulation of the primary visual cortex^8^, the lateral intraparietal area^9^, the projection from the frontal eye field to the superior colliculus^10^, and Purkinje cells in the cerebellum^26^ modulated saccadic eye movements in macaque monkeys. Optogenetic approaches were also applied to subcortical regions and modulated neuronal activity^27, 28, 29, 30^.

In the current study, we showed that oICMS in the M1 of macaque monkeys evoked clear forelimb movements that resembled movements evoked by eICMS (Figs. 3 and 4, Supplementary movie 1, Supplementary Table 2). The key to our success in inducing obvious movements by optogenetics in macaque monkeys was a winning combination of the following: (1) A highly effective AAV serotype DJ and a strong ubiquitous promoter CAG enabled potent gene transfer of ChR2 into neurons, (2) The AAV vector was injected into somatotopically identified regions of the M1, (3) Our ingenious optrode could identify layer 5 cortical neurons expressing ChR2 and illuminate both superficial and deep cortical layers expressing ChR2 by the optical fiber, and (4) Higher intensity of laser (51-1910 mW/mm^2^) was used for illumination than in previous studies (3-1600 mW/mm^2^) ^12, 13, 14^. Actually, oICMS changed neuronal activity more effectively (75% of cortical sites examined, Supplementary Table 2) than in previous reports (27.4-68.2% of neurons examined)^12, 13^.

AAV vectors are the most widely applied tools in optogenetics and have been successfully used to deliver opsins into the mouse, rat, and non-human primate brains. Previous studies evaluated transduction efficiency of different serotypes of AAV vectors, i.e., 1, 2, 5, 8, and 9, in the cortex of macaque monkeys^31, 32^, and reported low spreading and neuronal tropism of AAV2 and grater spreading of other serotypes. On the other hand, an AAV-DJ vector was created from DNA shuffling of eight AAV serotypes to mediate efficient gene transfer^33^ and showed high transduction effectiveness and neuronal tropism in the monkey putamen^34^. In the present study, in the macaque M1, we compared AAV2, AAV5, and AAV-DJ vectors expressing EGFP under the control of a ubiquitous CAG promoter that is commonly used in rodents. EGFP was successfully transduced by all AAV serotypes we injected, and the AAV-DJ vector showed the highest fluorescence intensity, the largest transduced areas and number of cells, and the highest protein expression (Supplementary Figs. 1, 2, 3). As an opsin for optical stimulation, we used a ChR2 variant, H134R, which was developed as a kinetic opsin variant in an early optogenetic study^4^ and obtained sufficient, obvious behavioral modulation, among many ChR variants developed, including kinetic opsin variants (ChETA, ChIEF)^35^, step-function opsins (C128S, D156A)^36, 37^, and red-shifted excitatory opsins (VChR1, Chrimson)^38, 39^. Thus, we prepared AAV-DJ vectors carrying CAG-hChR2(H134R)/EYFP or CAG-hChR2(H134R)/tdTomato to improve the expression levels of ChR2. In addition, we carefully performed electrophysiological mapping of the M1 and identified layer 5 of the forelimb region where large neuronal activity and eICMS evoked forelimb movements were clearly observed (Fig. 1a, b). We then injected the AAV vector into the identified region. Expression of hChR2/EYFP and hChR2/tdTomato was observed in the areas including layer 5 of the M1 (Fig. 1c).

In the present study, we used high intensity of laser (51-1910 mW/mm^2^) compared to previous studies. Previous studies targeting M1 cortical neurons expressing ChR2 used weaker laser (3-255 mW/mm^2^)^12, 13^ and could not affect behaviors. On the other hand, other studies used higher-intensity laser to inhibit Archaerhodopsin-T expressing superior collicular neurons (650-1600 mW/mm^2^)^14^ or to excite ChR2-expressing terminals in the superior colliculus (<1000 mW/mm^2^)^10^ and successfully affected eye movements. Thus, we decided to use higher-intensity laser in this study. Such intensity of laser seems to have no damage to the cortical neurons because (1) Cortical neurons could be activated repeatedly by oICMS after long oICMS without any injury discharges, (2) Neuronal and muscle activity could be induced by oICMS at the same site with the same threshold in different days, and (3) Histological examination showed no apparent damages in the cortex.

We compared body movements and EMG evoked by oICMS and by eICMS. Forelimb movements induced by oICMS were comparable to those induced by eICMS (Figs. 3 and 4, Supplementary movie 1). The latency of muscle activity evoked by oICMS (14.0±3.9 ms) was in the same range as that evoked by eICMS in the present (12.8±3.2 ms) and previous (8.8-10.1 ms)^40^ studies. Five-pulses oICMS induced similar amplitude of EMG as five-pulses eICMS did (Fig. 4), suggesting that both oICMS and eICMS activated the M1 to a similar extent. On the other hand, the area and peak amplitude of EMG evoked by one, two, three, and five pulses of oICMS and eICMS indicate that eICMS requires more pulses than oICMS to induce equivalent muscle activity (Fig. 4). These results suggest that oICMS induces body movements more effectively than eICMS. One reason is that oICMS illuminates neural tissue along the long axis and may effectively activate the cortical efferent zone^18, 41^. On the other hand, eICMS may activate neurons inside a sphere around the electrode tip, the size of which increases with increasing current^18, 19^: 10 µA and 100 µA currents activate neurons in a radius of 100 µm and 450 µm around the electrode tip. However, a recent study using two-photon calcium imaging indicated that eICMS may sparsely activate neurons around the electrode, sometimes as far as millimeters away^42^. Further studies are needed to elucidate the mechanism of oICMS especially in comparison with eICMS.

In this paper, we applied oICMS to the somatotopically identified region of the M1 and evoked forelimb movements successfully in a similar manner to eICMS. Our current study has removed obstacles to evoking movements by oICMS and opened a new window to the application of optogenetics to motor physiology in non-human primates: Selective manipulation of an optional population of neurons by using a region-specific promoter and/or pathway-specific labeling.

## Methods

### Animals

Six adult Japanese monkeys of both sexes (*Macaca fuscata*, body weight 3.0-10.5 kg) were used in this study (Supplementary Table 1). Experiments were performed at National Institute for Physiological Sciences (NIPS; Monkeys CL, NR, HJ, and PT) and Tohoku University (TU; Monkeys CH and HK). Each monkey was daily trained to sit quietly in a monkey chair. The experimental protocols were approved by the Institutional Animal Care and Use Committees of National Institutes of Natural Sciences and Tohoku University. All experiments were conducted according to the guidelines of the National Institutes of Health *Guide for the Care and Use of Laboratory Animals*.

### Surgery

Under general anesthesia with ketamine hydrochloride (10 mg/kg body weight, i.m.), xylazine hydrochloride (1 mg/kg, i.m.) and propofol (7 µg/kg, blood concentration) (NIPS) or with ketamine hydrochloride (10 mg/kg, i.m.), xylazine hydrochloride (0.5 mg/kg, i.m.) and isoflurane (1-1.5% in room air, 2 L/min) (TU), the monkeys underwent surgery to fix their heads painlessly to a stereotaxic frame that could be attached to a monkey chair as described previously^43, 44^. After skin incisions, the skull was widely exposed, and the periosteum was completely removed. Small polyether ether ketone (PEEK) screws were attached to the skull as anchors. The exposed skull and screws were completely covered with transparent acrylic resin. Two PEEK pipes were mounted in parallel over the frontal and occipital areas for head fixation. All surgical procedures were performed under aseptic conditions. Arterial oxygen saturation and heart rate were monitored during the surgery. Depth of anesthesia was assessed by heart rate and body movements. Additional anesthetic agents were administrated when necessary. Antibiotics and analgesics (ketoprofen) were administered (i.m.) after the surgery.

### Preparation of viral vectors

Adeno-associated virus (AAV) vectors were packaged using the AAV Helper Free Expression System (Cell Biolabs) as described previously^45^. The transfer plasmids, pAAV-CAG-EGFP and pAAV-CAG-hChR2(H134R)-EYFP, were constructed, and pAAV-CAG-hChR2(H134R)-tdTomato was obtained from Addgene. The packaging plasmids (pHelper and pAAV-2, pAAV-5 or pAAV-DJ) and transfer plasmid were co-transfected into HEK293T cells using the calcium phosphate method. Crude cell extracts containing AAV vector particles were purified by ultracentrifugation with CsCl, dialyzed, and concentrated with an Amicon 10K MWCO filter (Merck). The copy numbers of the viral genome (vg) determined by qPCR were 1.5×10^9^ vg/μl (CAG-EGFP), 4.0×10^9^ vg/μl (CAG-hChR2(H134R)/EYFP), and 4.3×10^9^ vg/μl (CAG-hChR2(H134R)/tdTomato), respectively.

### Mapping of the motor cortex and injection of viral vectors

After full recovery from the surgery, each monkey was seated quietly in a monkey chair with its head fixed in a stereotaxic frame. The skull over the central sulcus was removed under anesthesia with ketamine hydrochloride (10 mg/kg, i.m.) and xylazine hydrochloride (1 mg/ kg, i.m.) with local lidocaine or bupivacaine application. A rectangular plastic chamber was attached to cover the opening. The forelimb area of the primary motor cortex (M1) was identified by electrophysiological mapping^43, 44, 46, 47^ (Fig.1a, b; Supplementary Fig. 1a). Briefly, a glass-coated Elgiloy-alloy microelectrode was inserted perpendicularly to the cortical surface. Extracellular neuronal activity was recorded, and responses to somatosensory stimuli were examined. Then, intracortical microstimulation (ICMS; cathodal pulses, <50 µA strength, 0.2 ms pulse duration, 333 Hz, 12 pulses) was applied, and evoked movements were observed.

Based on the mapping, AAV vectors carrying the CAG-EGFP (to Monkeys CL and NR), CAG-hChR2(H134R)/EYFP (to Monkeys NR, HJ, and PT), or CAG-hChR2(H134R)/tdTomato (to Monkeys CH, HK, and NR) transgene were injected into the forelimb region of the M1 (Supplementary Table 1). A glass micropipette was made from a 3-mm glass capillary by using a puller (PE-2, Narishige) and connected to a 25-µl Hamilton microsyringe by Teflon tubing (JT-10, Eicom). A tungsten wire was inserted into a glass micropipette to record neuronal activity. The glass micropipette, tubing, and a Hamilton microsyringe were filled with Fluorinert (FC-3283, Sumitomo 3M). The syringe was mechanically controlled by a syringe pump (UMP3, WPI). Viral vector solution was loaded from the micropipette. The glass micropipette was inserted into the M1 perpendicularly to the cortical surface through a small incision in the dura mater. Injection sites were selected in layer 5 by recording large neuronal activity through the wire in the glass micropipette. Then, the vector solution (1 µl at each site) was injected slowly (50 nl/min). The micropipette was left in place for an additional 10 min and then slowly withdrawn. AAV vectors were injected into 2-5 sites (each serotype carrying CAG-EGFP, Supplementary Fig. 1a) or 4-15 sites (CAG-hChR2(H134R)/EYFP or CAG-hChR2(H134R)/tdTomato, Fig. 1a, b) in each hemisphere (Supplementary Table 1).

### Construction of optrodes

Optrodes with recording and stimulating electrodes were constructed from three major components (Fig. 1d, *left*): 1) a glass-coated tungsten electrode for recording (Alpha Omega), 2) an optical fiber with a 100-µm diameter silica core (CeramOptec) for optogenetic intracortical microstimulation (oICMS), and 3) a pair of 50-µm diameter tungsten wires for bipolar electrical intracortical microstimulation (eICMS), which were coated with Teflon except at their tips (California Fine Wire). The optical fiber was attached to the recording electrode 800-1000 µm behind the tip of the recording electrode. The pair of tungsten wires for eICMS was attached to the optical fiber (the distance between wire tips, 300-400 µm). The three components were fixed together by applying small drops of standard cyanoacrylate glue and epoxy glue, and then inserted into polyimide tubing (Microlumen). The base of the optorode was covered with stainless tubing (Fig. 1d, *right*). The tip of the optrode was observed under a digital microscope (VHX-1000 and 7000, Keyence) (Fig. 1d, *left*). The other end of the optical fiber was jacketed with a protective tubing with a length of around 15 cm and had a fiber-optic connector (Fig. 1d, *right*). The optical fiber was connected to a solid-state laser source (473 nm, 100 or 200 mW power output; COME2-473-100LS or COME2-LB473/200, Lucir). Light power was measured in mW using a power and energy meter (Ophir Optronics) at a distance of 1 mm from the light-emitting tips and calculated as power per unit area (mW/mm^2^).

### Optogenetic stimulation and extracellular recording in the M1

Three weeks after vector injection when ChR2 was expected to be expressed, recording and stimulating experiments were started. Each monkey was seated in a monkey chair under awake conditions. Two stainless rods were inserted into PEEK pipes over its head and were fixed in a stereotaxic frame without any pain as described previously^43, 44^. An optrode was mounted in a micromanipulator (MO-81-S, Narishige) and inserted into the M1 perpendicularly to the cortical surface through a small incision of the dura mater. Electrophysiological signals from the recording electrode were amplified (5000 times), filtered (300-3000 Hz), digitized at 50 kHz, and stored on a computer using LabView software (National Instrument). For oICMS, the optical fiber was connected to the solid-state laser source, which was controlled by a stimulator (SEN8201, Nihonkohden). Single-pulse (0.4-15 mW strength corresponding to 51-1910 mW/mm^2^, 0.1-10 ms pulse duration) or repetitive (0.4-15 mW strength, 0.1-2 ms pulse duration, 100-333 Hz, 2-50 pulses) oICMS was applied through the optical fiber at every 1.4 or 2.4 s. The stimulus frequency of repetitive oICMS followed that was commonly used in repetitive eICMS. For eICMS, the tungsten wires for bipolar stimulation were connected to an isolator (SS-202J, Nihonkohden) and a stimulator (SEN8201), and single-pulse (monophasic, 20-50 µA strength, 0.2 ms pulse duration) or repetitive (monophasic, 20-65 µA strength, 0.2 ms pulse duration, 100-333 Hz, 2-20 pulses) eICMS was applied through the bipolar stimulating electrodes at every 1.4 s.

### EMG recording

Body movements and electromyogram (EMG) were observed when monkeys were at rest without any movements. Body movements evoked by eICMS or oICMS were video-recorded. The timing of oICMS was indicated by the flash of LEDs set near the body parts recorded. When body movements were observed, a pair of surface dish electrodes (NE-05, Nihonkohden) were attached to the skin over the belly of the responsible muscle. Muscle activity was amplified (5000 times), filtered (50-3000 Hz), digitized at 50 kHz, and stored on a computer simultaneously with neuronal activity using LabView software. Although body parts in which movements and/or EMG were evoked depended on the cortical site of oICMS/eICMS, EMG of the extensor carpi radialis longus and flexor carpi radialis muscles was mainly recorded and used for further analyses.

### Histological examination

After collection of enough data (in the case of electrophysiological experiments) or 2-4 weeks after AAV injections (for evaluation of AAV vectors), monkeys were deeply anesthetized with sodium pentobarbital (50 mg/kg, i.v.) and perfused transcardially with 0.1 M phosphate buffer (pH 7.3) followed by 10% formalin in 0.1 M phosphate buffer, and 0.1 M phosphate buffer containing 10% sucrose. The brains were removed and kept in the same buffer containing 30% sucrose at 4 °C. Frontal sections (50 μm thick) were cut serially with a freezing microtome and collected in 0.01M PBS. Every sixth sections were placed onto gelatine-coated glass slides, air dried, and mounted in Vectashield (Vector Laboratories). The remaining sections were used for immunohistochemistry to enhance fluorescence. Every sixth free-floating sections were incubated with primary antibodies against rabbit GFP (1:1000; Thermo Fisher Scientific) or rabbit DsRed (1:500; Takara Clontech) at 4°C overnight, and then visualized with secondary antibodies conjugated with Alexa Fluor 488 or Alexa Fluor 594. All fluorescent images were captured by a fluorescence microscope (BZ-X710 and BZ-X Viewer, Keyence).

### Data and statistical analyses

Recorded EMG was rectified, re-sampled at 2 kHz, and averaged for 30 stimulus trials using Igor software (WaveMetrics). The mean and SD values of EMG during the 100-ms period preceding the onset of stimulation were considered as the baseline activity. Changes in EMG in response to stimulation were judged significant if the EMG during at least six consecutive points (3 ms) reached a significant level of *p*<0.01 (one-tailed *t* test). The latency was defined as the time at which the first point exceeded this level. The responses were judged to end when six consecutive points fell below the significance level. The end point of the responses was determined as the time at which the last point exceeded this level. The area of EMG was defined as the area of averaged EMG during the significant response over the baseline. The peak amplitude of EMG was defined as the height of maximal peak in the averaged EMG. Data containing weak EMG, defined as small area (<20 mV•ms) or amplitude (<2 mV) of EMG evoked by oICMS or eICMS (five pulses at 200 Hz) were excluded from further analyses. The area and peak amplitude of EMG were compared between different stimulus pulse numbers and between different stimulus frequencies by Friedman test with Dunn’s post hoc test. Significant level was set to *p*<0.05. Statistical analyses were performed using Prism software (GraphPad).

## Acknowledgements

We thank S. Sato, H. Isogai, N. Suzuki, K. Awamura, K. Miyamoto, T. Sugiyama, R. Kageyama, M. Takahashi, Y. Kajita, and the Equipment Development Center (Institute for Molecular Science) for technical support and Y. Yamagata for her critical reading of the manuscript. This work was supported by MEXT KAKENHI (“Non-linear Neuro-oscillology”, 15H05873 to A.N. and 15H05879 to H.M., and “Adaptive Circuit Shift”, 17H05590 to H.S.), JSPS KAKENHI (15K01854 to H.W., 16K01968 to H.S., 16K07014 to S.C., 26290001 to H.M., and 26250009 to A.N.), JST CREST (JPMJCR1853 to S.C.), and AMED (19dm0307005h0002 and 19dm0207050h003 to A.N.). Four Japanese monkeys used in the present study were provided through National Bio-Resource Project (NBRP) “Japanese Monkeys” of AMED.

## Author contributions

H.W., H.S., S.C., H.M., and A.N. designed the study. H.S. and K.K. generated the viral vectors. Y.F. and M.F. measured the expression of viral vectors. H.W., H.S., S.C., and A.N. performed the experiments. H.W., H.S., and S.C. analyzed the data. H.S., S.C., and A.N. wrote the manuscript.

## Data accessibility

The data presented in the current manuscript are available upon request to the corresponding author.

