## Supplementary Data for "Forelimb movements evoked by optogenetic stimulation of the macaque motor cortex"

Watanabe et al.

List of Contents

Supplementary Methods

Supplementary Tables 1 and 2

Supplementary Figures 1-5

**Supplementary Methods**

**Histological examination and analysis.** Every sixth free-floating sections of the monkey brain were incubated with primary antibodies against rabbit GFP (1:1000; Thermo Fisher Scientific), mouse NeuN (1:400; Merck), mouse parvalbumin (PV) (1:5000, Swant), and mouse GFAP (1:400; Sigma) at 4℃ overnight, and then visualized with secondary antibodies conjugated with Alexa Fluor 488 or Alexa Fluor 594 (1: 1000, Thermo Fisher Scientific). All fluorescent images were captured by a fluorescence microscope (BZ-X710 and BZ-X Viewer, Keyence). For measuring the extent of the areas covered by EGFP positive cells, hChR2(H134R)/EYFP or hChR2(H134R)/tdTomato signals were manually surrounded and measured using ImageJ software (National Institutes of Health). For measuring the fluorescence intensity of EGFP, all fluorescent images were captured with same exposure, then individual EGFP positive cells in 500 μm × 500 μm region of interest (ROI) were surrounded manually, and the optical density was measured using ImageJ software in 8-12 ROIs. The fluorescence intensity by AAV2 injection was averaged over all cells and used as the reference value. The fluorescence intensity by AAV2, AAV5, and AAV-DJ injection was expressed as a ratio to the reference value. For counting of EGFP, NeuN, PV, and GFAP positive cells, 10-55 ROIs (500 μm X 500 μm) were examined manually using ImageJ software.

**Quantitative western blotting.** To compare AAV2, AAV5, and AAV-DJ vectors, we injected one of these vectors into the mouse brain and performed western blot analyses. Experiments were performed at National Institute for Physiological Sciences. The experimental protocols were approved by the Institutional Animal Care and Use Committees of National Institutes of Natural Sciences. AAV2, AAV5, or AAV-DJ vector carrying the CAG-EGFP transgene ($4\times10^9$ viral genome (vg)/μl, 0.5 μl/site, two sites) was injected into each hemisphere of the mouse under general anesthesia with isoflurane (1.0-1.5%). Two weeks after AAV injection, mice were sacrificed by cervical dislocation, then the brain hemispheres were dissected and homogenized separately with 10-volume of buffer containing 20 mM Tris-HCl (pH 8.0), 1 mM EDTA, 320 mM sucrose, and 100 µg $mL^{-1}$ phenylmethylsulfonyl fluoride. The protein concentration of individual homogenates was measured by BCA protein assay (Thermo Fisher Scientific), and the homogenates were diluted to 1 mg/ml in SDS-PAGE sample buffer. Individual homogenates (15 µg each) were subjected to western blotting with anti-GFP and anti-β-catenin (as an internal control) antibodies. Anti-GFP rabbit polyclonal antibody was raised against GST-GFP and affinity-purified. Anti-β-catenin mouse monoclonal antibody (clone, 14/beta-catenin) was purchased from BD Biosciences. For quantitative western blotting, chemical luminescent signals were detected with the FUSION Solo system (Vilber-Lourmat) and analyzed with the FUSION software.

### Supplementary Tables

| Monkey | Male/ female | BW (kg) | | Viral vector(s) injected | Total volume injected (μl) | Experiments | Survival period |
|---|---|---|---|---|---|---|---|
| CL | Female | 3.0 | Left M1 | AAV2-CAG-EGFP | 2 | Histology | 1 m |
| | | | | AAV5-CAG-EGFP | 2 | Histology | 1 m |
| | | | | AAV-DJ-CAG-EGFP | 3 | Histology | 1 m |
| CH | Female | 6.7 | Left M1 | AAV-DJ-CAG-H134R-tdTomato | 8 | oICMS/Histology | 6 m |
| HK | Male | 10.5 | Left M1 | AAV-DJ-CAG-H134R-tdTomato | 8 | oICMS/Histology | 6 m |
| NR | Female | 6.5 | Left M1 | AAV-DJ-CAG-H134R-EYFP | 4 | Histology | 2 w |
| | | | | AAV2-CAG-EGFP | 4 | Histology | 2 w |
| | | | | AAV5-CAG-EGFP | 4 | Histology | 2 w |
| | | | | AAV-DJ-CAG-EGFP | 5 | Histology | 2 w |
| | | | Right M1 | AAV-DJ-CAG-H134R-EYFP | 11 | oICMS/Histology | 2.5 m |
| | | | | AAV-DJ-CAG-H134R-tdTomato | 4 | Histology | 5 w |
| HJ | Female | 7.5 | Right M1 | AAV-DJ-CAG-H134R-EYFP | 15 | oICMS/Histology | 5 m |
| PT | Female | 6.1 | Left M1 | AAV-DJ-CAG-H134R-EYFP | 14 | oICMS | * |

**Supplementary Table 1.** Summary of experiments. Survival period, period in months (m) or weeks (w) between AAV injection and sacrifice.

* Monkey PT is still alive and has not been histologically verified yet.

| Monkey | oICMS | | | | eICMS | |
|---|---|---|---|---|---|---|
| | Neuronal activity (evoked sites/examined) | No of activated neurons | Movements (evoked sites/examined) | Muscle activity (evoked sites/examined) | Movements (evoked sites/examined) | Muscle activity (evoked sites/examined) |
| CH | 17/19 | 29 | 7/19 | 0/0 | 0/0 | 0/0 |
| HK | 44/60 | 88 | 7/60 | 5/10 | 0/0 | 0/0 |
| NR | 62/85 | 109 | 51/85 | 48/80 | 2/2 | 2/2 |
| HJ | 45/55 | 85 | 43/55 | 48/53 | 14/16 | 14/16 |
| PT | 22/35 | 48 | 24/25 | 23/24 | 15/18 | 15/18 |
| Total | 190/254 | 359 | 132/244 | 124/167 | 31/36 | 31/36 |
| % | 75% | | 54% | 74% | 86% | 86% |

**Supplementary Table 2.** Summary of optogenetic and electrical intracortical microstimulation. Numbers of sites where optogenetic intracortical microstimulation (oICMS) evoked neuronal activity, movements, and muscle activity among numbers of sites examined, numbers of neurons activated by oICMS, and numbers of sites where electrical intracortical microstimulation (eICMS) evoked movements and muscle activity among numbers of sites examined, are shown. Numbers of activated neurons include multiunit activity, which was sometimes induced by oICMS.

**Supplementary Figures**

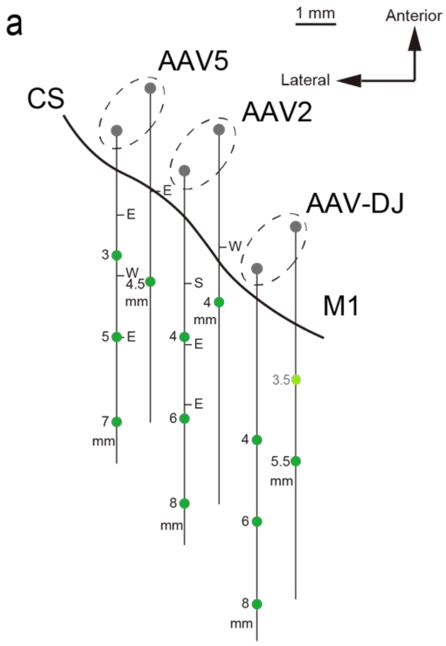


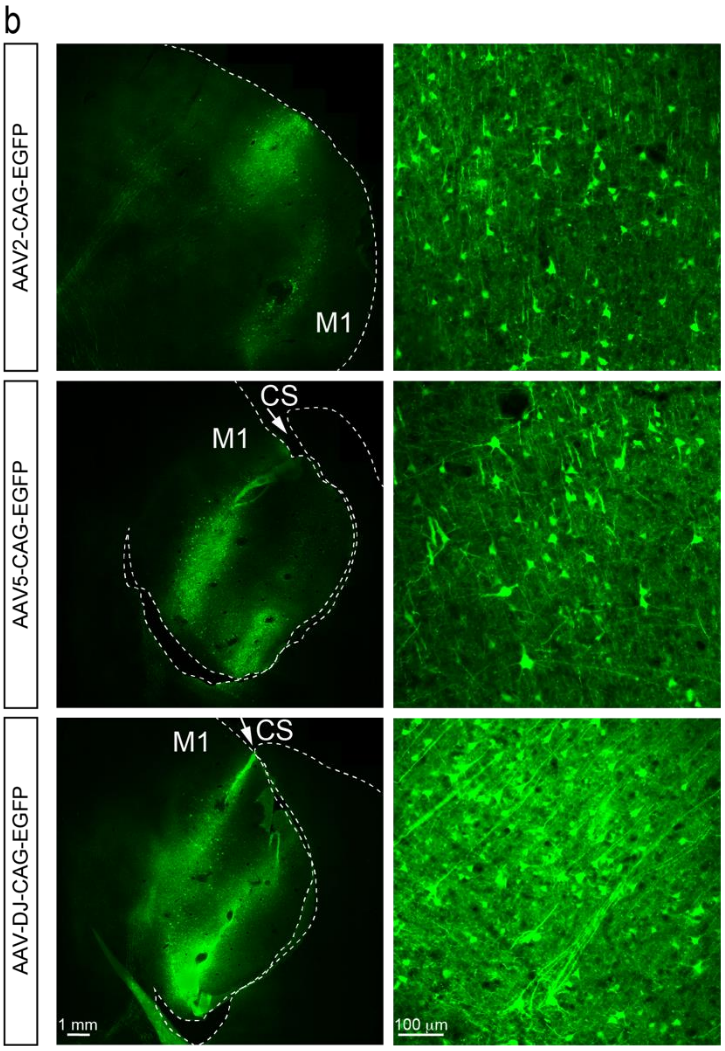


c

| | Area ($mm^2$) | Density (cells / ROI) |
|---|---|---|
| AAV2 | 17.4 | 87.9 |
| AAV5 | 44.2 | 66.8 |
| AAV-DJ | 70.6 | 99.7 |

**Supplementary Figure 1** Expression of EGFP transduced by AAV2, AAV5, and AAV-DJ vectors. **a** Injection sites in the primary motor cortex (M1). AAV2, AAV5, and AAV-DJ vectors were injected at different depths from the cortical surface (indicated by the green circles) in the anterior bank of the central sulcus (CS) of Monkey NR. Each letter indicates a somatic body part: E, elbow; S shoulder; W wrist. Histological data at four sites of each vector injection were analyzed. The data at the light green circle by AAV-DJ injection was excluded from histological analysis. **b** Expression of EGFP mediated by AAV2, AAV5, and AAV-DJ vectors in the M1 shown in frontal sections (Monkey NR). The transduced areas and the intensity of fluorescence signals were more prominent around the injection sites of the AAV-DJ vector, compared with other serotypes. Neuropil as well as somata express strong fluorescence by AAV-DJ injection (*bottom right*). **c** Areas covered by EGFP positive cells (Monkey NR) and number of EGFP positive cells (Monkey CL and NR) per ROI (500 µm × 500 µm). EGFP positive cells were observed in 8, 10, and 12 slices after AAV2, AAV5, and AAV-DJ injections, respectively, in Monkey NR.

a

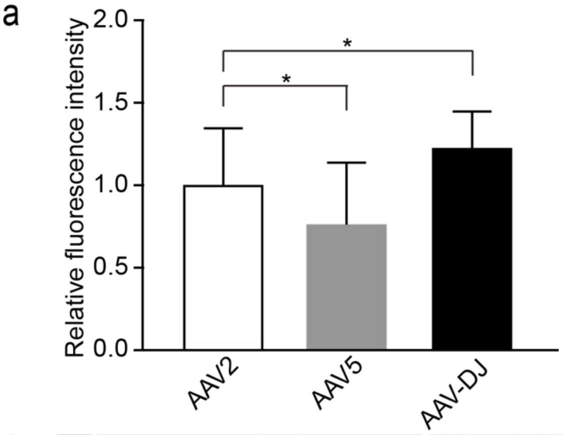


b

| | NeuN / EGFP (%) | PV / EGFP (%) | GFAP / EGFP (%) |
|---|---|---|---|
| AAV2 | 88.3 | 4.6 | 1.8 |
| AAV5 | 88.7 | 8.5 | 6.8 |
| AAV-DJ | 91.2 | 15.9 | 1.9 |

c

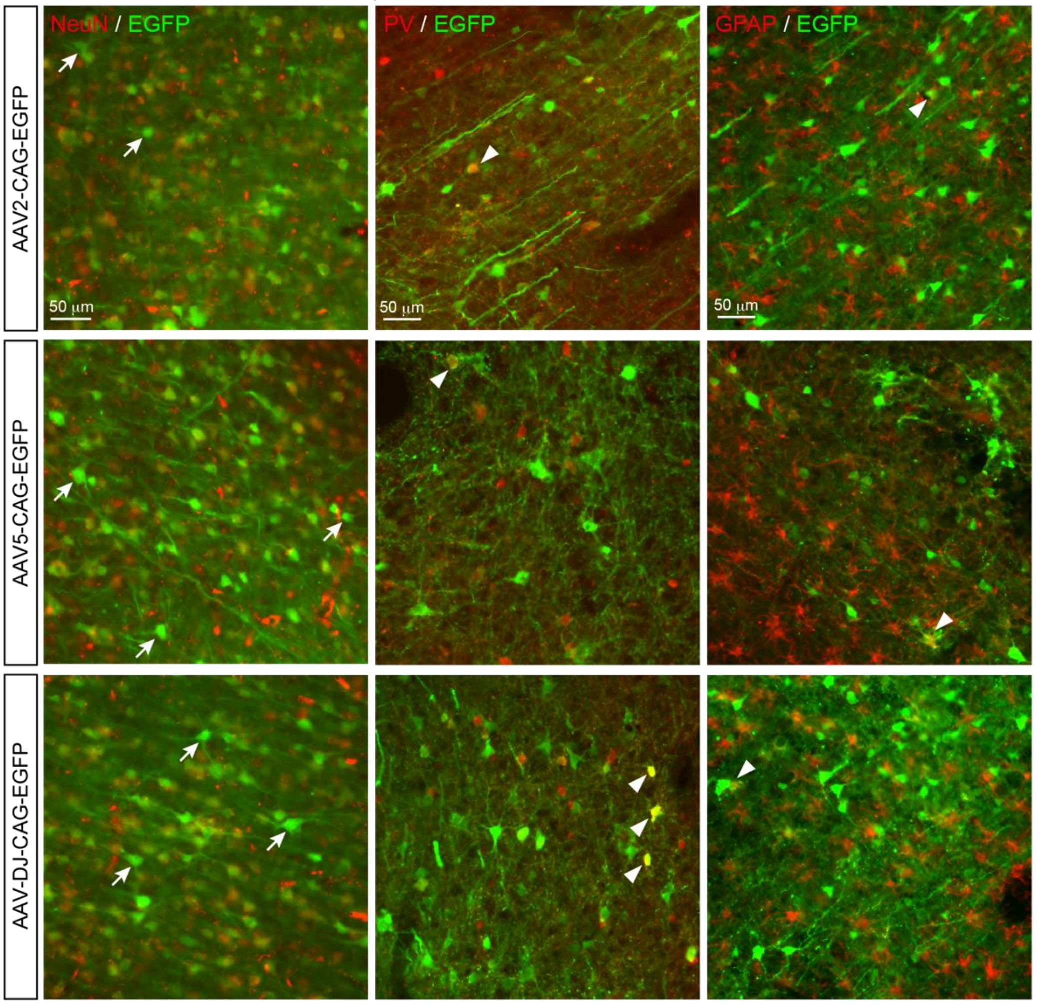


d

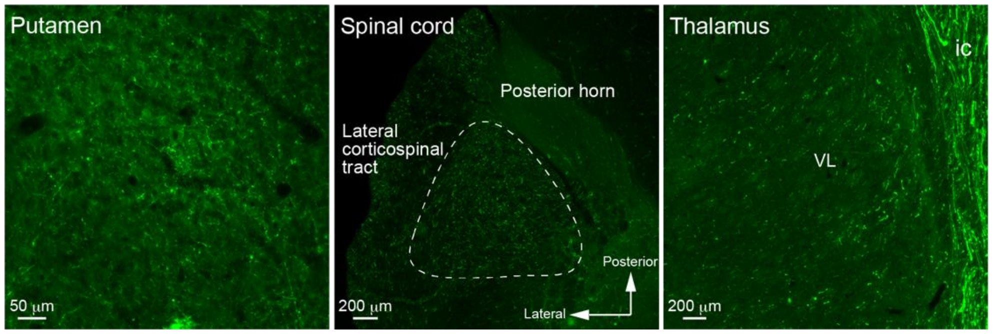

**Supplementary Figure 2** Characterization of EGFP positive cells transduced by AAV2, AAV5, and AAV-DJ vectors. **a** The relative fluorescence intensity (mean and SD) per cell by AAV2, AAV5, and AAV-DJ injection expressed as a ratio to the reference value by AAV2 injection in Monkey NR (fluorescence intensities of AAV2 averaged over all cells). *$p<0.05$, one-way ANOVA followed by Tukey's post hoc test. **b** Percentages of NeuN (marker for neuron), palvalbmine (PV, marker for PV containing interneurons), and GFAP (markers for glial cells) positive cells among EGFP positive cells. NeuN, PV, and GFAP were examined in Monkeys CL, NR, and NR, respectively. **c** Double staining with anti-NeuN (*left column,* Monkey CL), anti-PV (*middle column*, Monkey NR) or anti-GFAP (*right column,* Monkey NR) antibodies in combination with anti-GFP antibodies. Most EGFP-positive cells are NeuN-positive, and EGFP single-positive cells is minor (arrows). On the hand, a small population of neurons are PV/EGFP or GFAP/EGFP double-positive (arrow heads). **d** EGFP expression in the putamen (*left*), spinal cord at the level of C7 (*middle*), and ventrolateral nucleus (VL) of the thalamus (*right*) in Monkey CL. Labeled terminals in the putamen and labeled axons in the lateral funiculus of the spinal cord, presumably the lateral corticospinal tract (circled with broken lines) were observed. No retrogradely EGFP-labeled cells were found in the VL. ic, internal capsule.

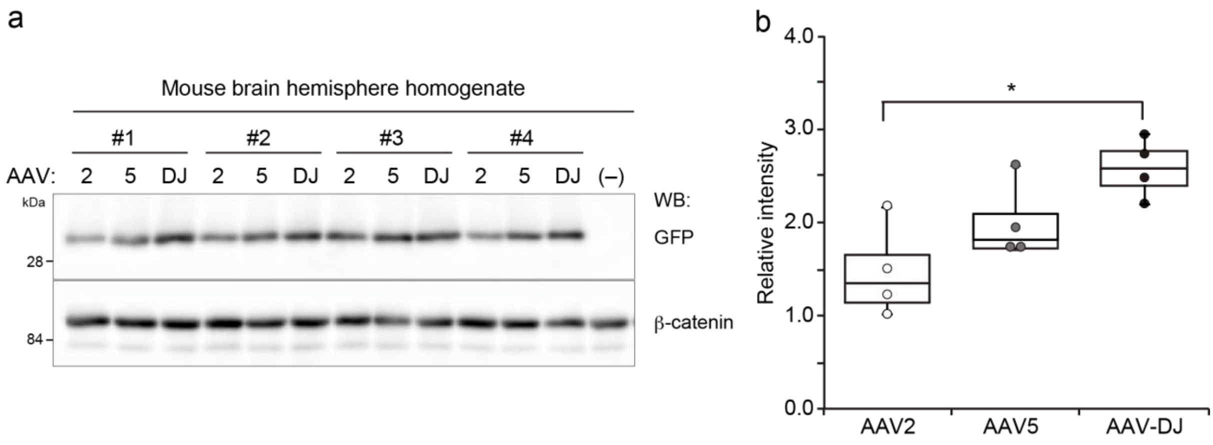


**Supplementary Figure 3** Quantitative analysis of EGFP expression transduced by AAV2, AAV5, and AAV-DJ vectors. **a** Quantitative western blotting of brain homogenates with anti-GFP and anti-β-catenin (control) antibodies. AAV2, AAV5, or AAV-DJ vector was separately injected to both hemispheres of two mice (n=4 for each vector). The AAV-DJ vector induced higher expression of EGFP than AAV2 and AAV5. **b** Box plots showing the median (center line), upper and lower quartiles (box), and 1.5 × interquartile range from upper and lower quartiles (whiskers) of the relative intensities in AAV5 and AAV-DJ vectors as a ratio to that in experiment #1 of AAV2. *$p<0.05$, one-way ANOVA followed by Tukey's post hoc test.

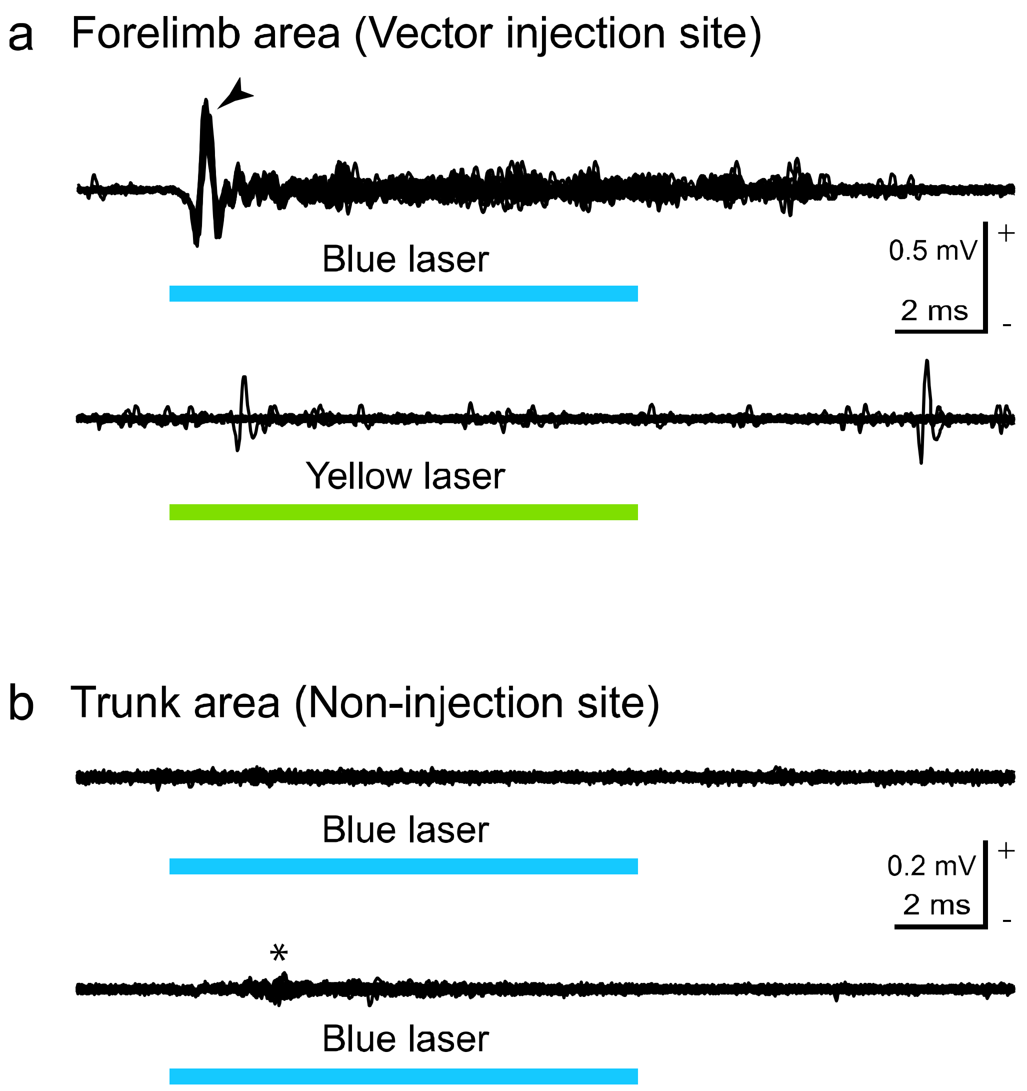


**Supplementary Figure 4** Electrical potentials evoked by oICMS. **a** Electrical potentials recorded at a vector injection site (forelimb area of the M1). oICMS with 473 nm blue laser (15 mW corresponding to 1910 mW/mm$^2$, 10 ms, single pulse) induced a large deflection (arrowhead in *upper trace*), probably composed of action potentials with a short and constant latency, followed by small spikes. On the other hand, oICMS with 589 nm yellow laser (15 mW, 10 ms, single pulse; 100mW power output, COME2-589-100LS, Lucir) delivered through the same optical fiber did not induce any responses at the same recording site (*lower trace*). **b** Electrical potentials recorded at a non-injection site (trunk area of the M1). oICMS with 473 nm blue laser (15 mW, 10 ms, single pulse) induced no responses (*upper trace*) or only small fluctuations of the baseline (* in *lower trace*), probably synaptically induced by excited axons of ChR2 expressing cortical neurons. These observations suggest that the large deflection observed in the upper trace in **a** is derived from neuronal activity, but not from a nonspecific photoelectric effect.

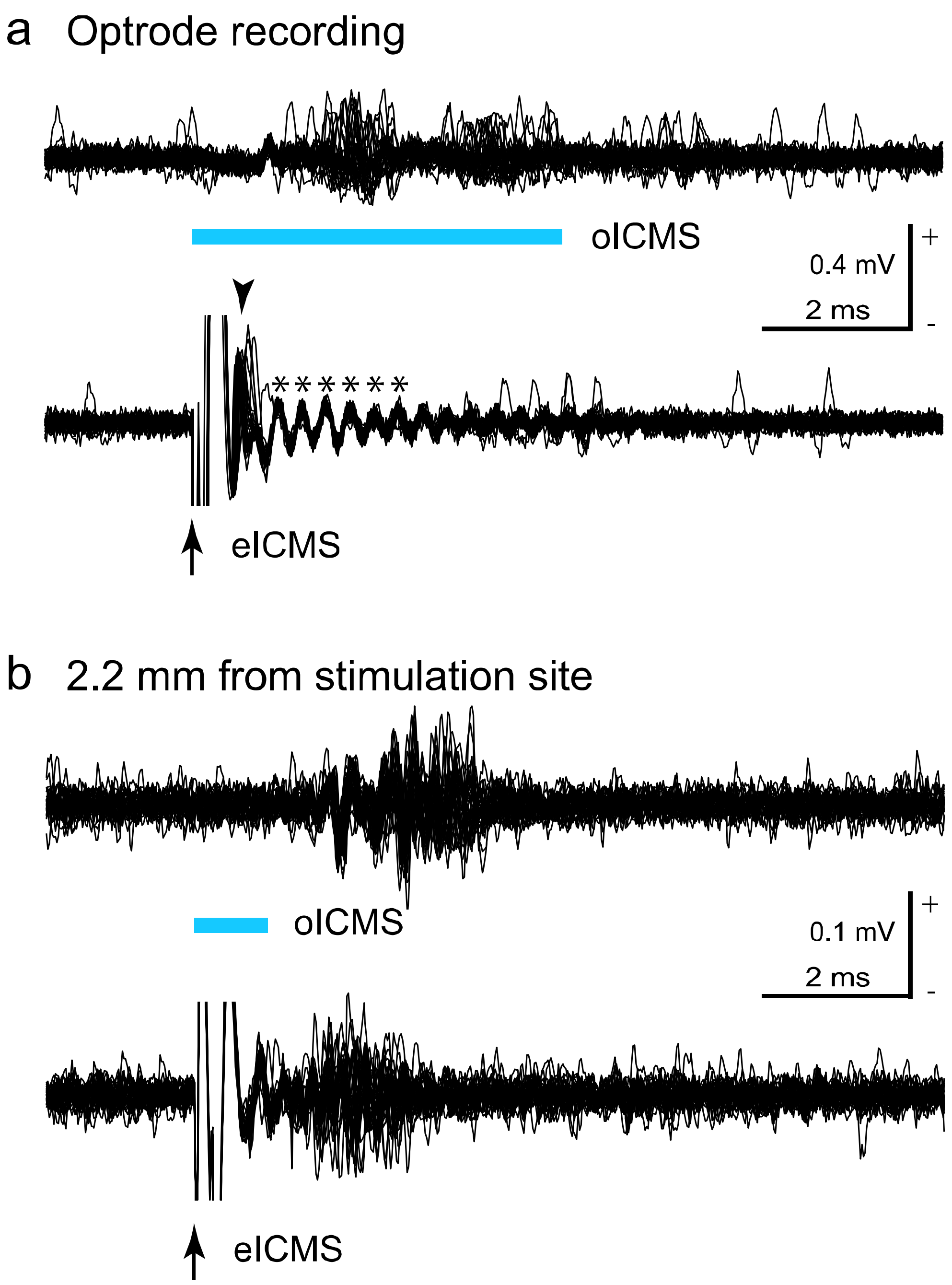


**Supplementary Figure 5** Neuronal activity evoked by oICMS and eICMS. **a** Neuronal activity recorded using an optrode. oICMS (15 mW corresponding to 1910 mW/mm$^2$, 5 ms, single pulse) induced small spikes (*upper trace*), while eICMS (65 µA, 0.2 ms, single pulse) evoked a large spike (arrowhead) followed by oscillatory activity (*) at the same recording site (*lower trace*). **b** Neuronal activity recorded at a horizontal distance of 2.2 mm from the oICMS/eICMS site using another recording electrode. Both oICMS and eICMS induced spikes in a similar group of neurons with larger time jitter at longer latencies than those observed in **a**, suggesting that they were induced transsynaptically by inputs from excited neurons at the oICMS/eICMS site.
